# Dynamics of the COVID -19 Related Publications

**DOI:** 10.1101/2020.08.05.237313

**Authors:** Biplab Sarkar, Anusheel Munshi, Bhaswar Ghosh, Tharmarnadar Ganesh, Arjunan Manikandan, Subhra Singhdha Biswal, Tanweer Sahid, B Rajagopalan, Sinjini Snegupta, Sandipan Raychaudhuri, Jibak Bhattacharya, Mahasin Gazi, Arundhati De, Kirubha George, Tanmay Ghosh, Jawed Akhtar, Sourav Mondal, Mukti Mukherjee, Rosalima Gupta, Soumya Roy, Kanan Jassal, Khushboo Rastogi, Kanika Bansal, Prasenjit Chatterjee, Litan Naha Biswas, Shymasish Bondopadhay, Anirudh Pradhan, Bidhu Kalyan Mohanti

## Abstract

**Background:** This study aims to analyze the dynamics of the published articles and preprints of Covid-19 related literature from different scientific databases and sharing platforms.

**Methods:** The PubMed, Elsevier, and Research Gate (RG) databases were under consideration in this study over a specific time. Analyses were carried out on the number of publications as (a) function of time (day), (b) journals and (c) authors. Doubling time of the number of publications was analyzed for PubMed “all articles” and Elsevier published articles. Analyzed databases were (1A) PubMed “all articles” (01/12/2019-12/06/2020) (1B) PubMed Review articles (01/12/2019-2/5/2020) and (1C) PubMed Clinical Trials (01/01/2020-30/06/2020) (2) Elsevier all publications (01/12/2019-25/05/2020) (3) RG (Article, Pre Print, Technical Report) (15/04/2020–30/4/2020).

**Findings:** Total publications in the observation period for PubMed, Elsevier, and RG were 23000, 5898 and 5393 respectively. The average number of publications/day for PubMed, Elsevier and RG were 70.0 ±128.6, 77.6±125.3 and 255.6±205.8 respectively. PubMed shows an avalanche in the number of publication around May 10, number of publications jumped from 6.0±8.4/day to 282.5±110.3/day. The average doubling time for PubMed, Elsevier, and RG was 10.3±4 days, 20.6 days, and 2.3±2.0 days respectively. In PubMed average articles/journal was 5.2±10.3 and top 20 authors representing 935 articles are of Chinese descent. The average number of publications per author for PubMed, Elsevier, and RG was 1.2±1.4, 1.3±0.9, and 1.1±0.4 respectively. Subgroup analysis, PubMed review articles mean and median review time for each article were <0|17±17|77> and 13.9 days respectively; and reducing at a rate of-0.21 days (count)/day.

**Interpretation:** Although the disease has been known for around 6 months, the number of publications related to the Covid-19 until now is huge and growing very fast with time. It is essential to rationalize the publications scientifically by the researchers, authors, reviewers, and publishing houses.

**Funding:** None

## Introduction

On March 11, 2020, WHO declared COVID-19 as a global pandemic. This virus rapidly crossed borders and led to a major healthcare crisis and economic slowdown.

Most international health organizations have stated an urgent need to stop, control and reduce the impact of the virus at every opportunity (1).

Healthcare systems, various sectors of industry, and overall economy of the globe have been hit severely by this pandemic. As of July 23, 2020, there were 15 million confirmed cases worldwide with the number of fatalities in excess of 600, 000 so far. [1]

How has the scientific world reacted to this pandemic? This pandemic has brought to fore the gaping incoherence in opinions expressed by agencies the world over. Government bodies, non-government institutions, pharmaceutical industry, researchers have all made their sounds, but without much unison.

A sudden outbreak in the number of publications has been observed in the last few months. By one estimate, the COVID-19 literature published since January has reached more than 23,000 papers and is doubling every 20 days—among the biggest explosions of scientific literature ever. [2]

Another study says the covid-19 literature doubling time is 14 days. A quick run of the total count of publications with the keyword “COVID-19”, on Google Scholar shows about 20,500 results for 2019-2020. For the same period PubMed shows 23480 for the keyword “COVID-19” [All Fields] [as on 12-06-2020]. In addition to the staggering numbers, the pandemic has also affected the quality and content of publications. Editors across the world have gone to the extent of appointing “COVID Editors”, who supposedly are to take care of the section of the journals that deals with COVID articles. Professional bodies, both national and international, of every specialty have attempted to bring out their own “guidelines” and “recommendations” to deal with-patients of their specialty in the “COVID situation”. Researches seem to write articles endlessly, often repeating and restating what is already written.

This also raises questions on the quality of data and the authenticity of the results. On April 29, 2020, after a hurriedly conducted trial, the National Institute of Allergy and Infectious Diseases proposed Remdesivir, as an effective drug for this virus. [3] A contrary opinion was given in an earlier article published in a reputed journal. This trial showed no benefit of using Remdesivir. [4] The use of chloroquine is another example that stands out as an example, with divergent opinions from all quarters.[5]

The need of the hour is to print and publish only credible, proven information on respective platforms, including online and printed journals. The flooding of irrelevant junk needs to be stopped. COVID pandemic came and hopefully should be over soon, but this irrelevant information junk shall remain stored online for all times to come.

This study aims to evaluate the characteristics of the authors and topics from different databases and publishing houses. In a subgroup analysis, nationalities of the corresponding authors were analyzed.

## Materials and methods

Different medical databases, publication houses and sharing platforms were analyzed, each over a certain period of time to obtain the different characteristics and dynamics of the published/uploaded articles. All the databases created a separate section to tackle the Covid-19 related literature. The keywords in PubMed were “COVID-19” OR “Coronavirus” OR “Corona virus” OR “Coronaviruses”. [6] Elsevier and Research Gate (RG) were having a separate database for Covid-19 specific research. [7–8]

The data for different databases were extracted as a spreadsheet (PubMed) or saved first as a html (Elsevier and RG) file which was then converted into a spreadsheet. Several articles and write-ups with only meager relationship to the disease were not further counted in the analysis.

Analyzed databases were (1a) PubMed “all articles” (12/06/2020 - 01/12/2019), (1b) PubMed review articles (02/05/2020-01/12/19), with a subgroup analysis of total review time, (1c) PubMed clinical trials (January-June 2020); (2) Elsevier all publications except Erratum (December 2019-25-05-2020) (3) Research Gate (Published article, Pre-print, Technical Report) (15/04/2020 - 30/04/2020).

The dynamics of covid-19 related documents in PubMed, Elsevier, Research Gate databases were analyzed for number of publications as (a) function of time (day), (b) journals and (c) authors. Number of publications as a function of doubling time was analyzed for PubMed “all articles” and Elsevier published articles.

Analysis of total review time (submission date to acceptance date) was done for 150 review articles published in Elsevier and Covid-19 related control trials presented in PubMed.

Average values quoted in this article as <Minimum Value | Average ± Standard Deviation | Maximum value>, where maximum-minimum indicate the range.

## Results

### PubMed: “all articles”

### Number of Publications Vs Days

Total number of publications in PubMed: “all articles” was 23480. Analysis was limited to the last 10000 articles. Figure-1a shows the number of publications until 13/06/2020 as a function of date and a 5-days moving average. The overall average (± Standard Deviation) number of publications is 70.0 ±128.6 /day. The avalanche on number of publications that occurred after May 10, 2020 is seen from a look at the number of average daily publications that jumped significantly from 6.0±8.4 to 282.5±110.3 (figure-1b). As per the characteristics of the cumulative number, the publications were divided into two groups: from 20/02/2020 to 10/05/2020 (Group-1) and from 11/05/2020 to 13/06/2020 (Group-2). In Group-1, the cumulative number of publications (figure-2a) displayed a parabolic relationship against the days with a major coefficient a=0.09. In the post-avalanche Group-2, it changed to a linear relationship with a slope of m= 280.2 pointing to the average number of publications per day (282.5) during that period. Post-avalanche increase in the number of publications per day was 4670%.

**Figure-1a:**
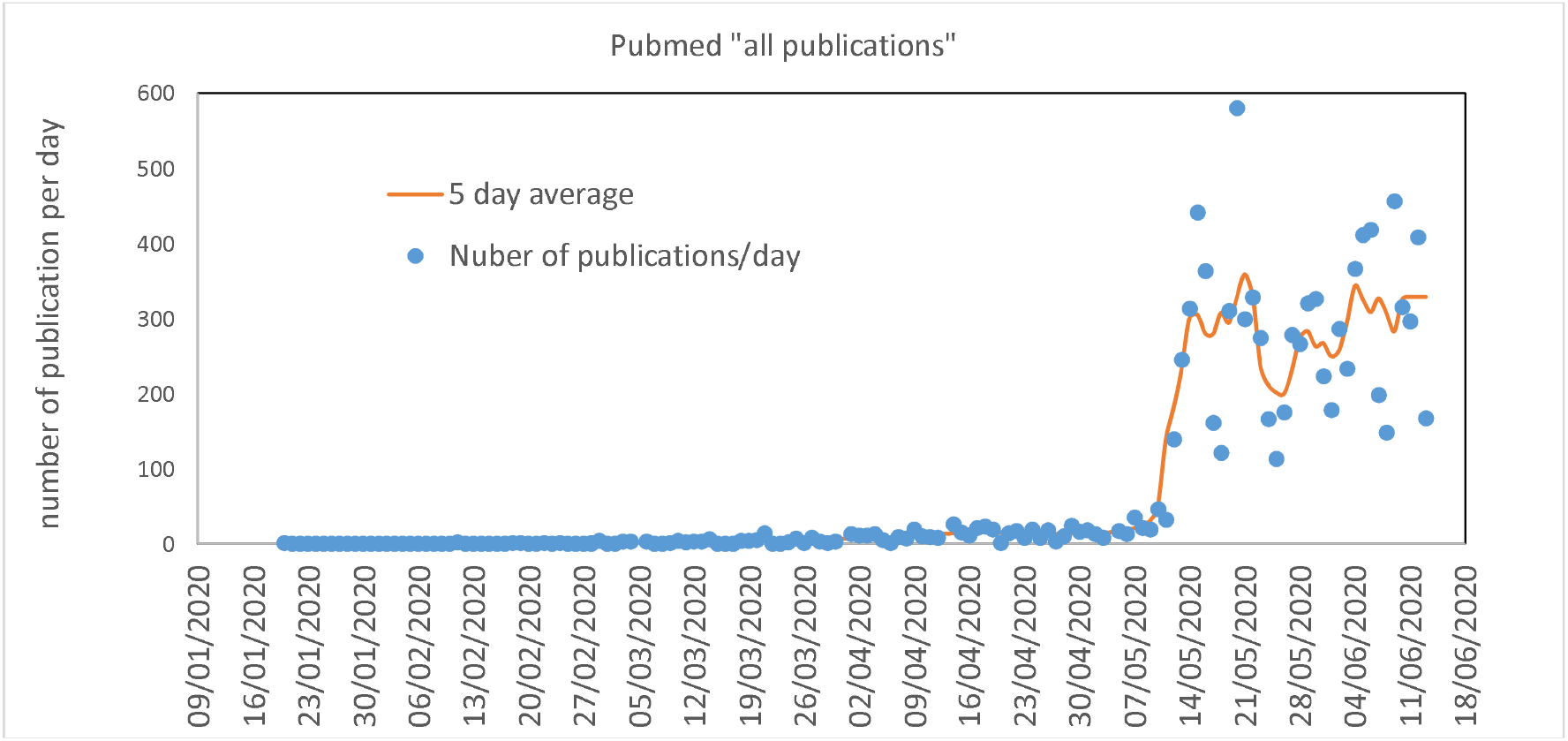
Publication as a function of date for all articles presented in PubMed

**Figure-1b:**
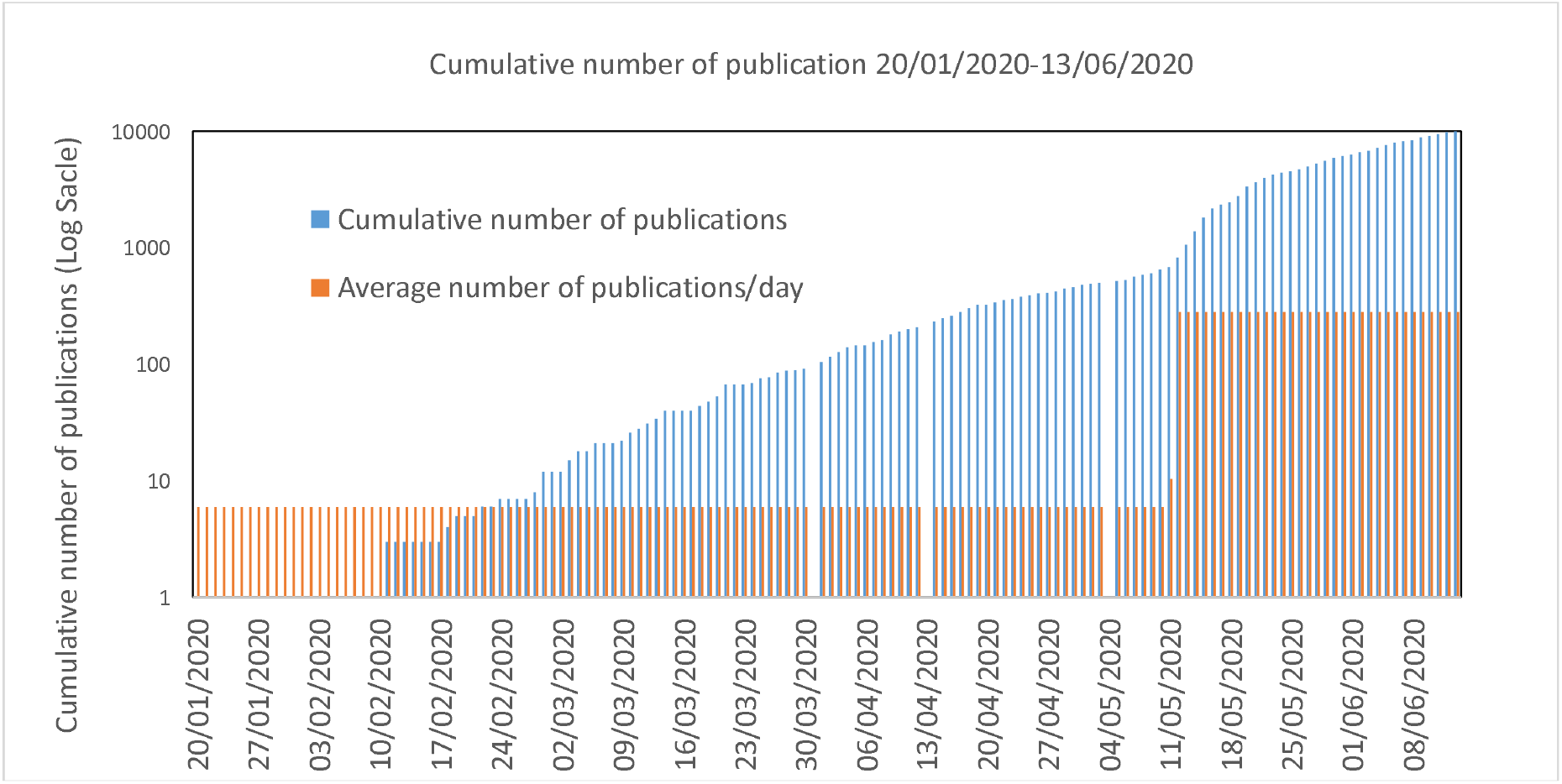
Cumulative and average number of publications 20/01/2020-13/06/2020 as a function of date

**Figure-2a:**
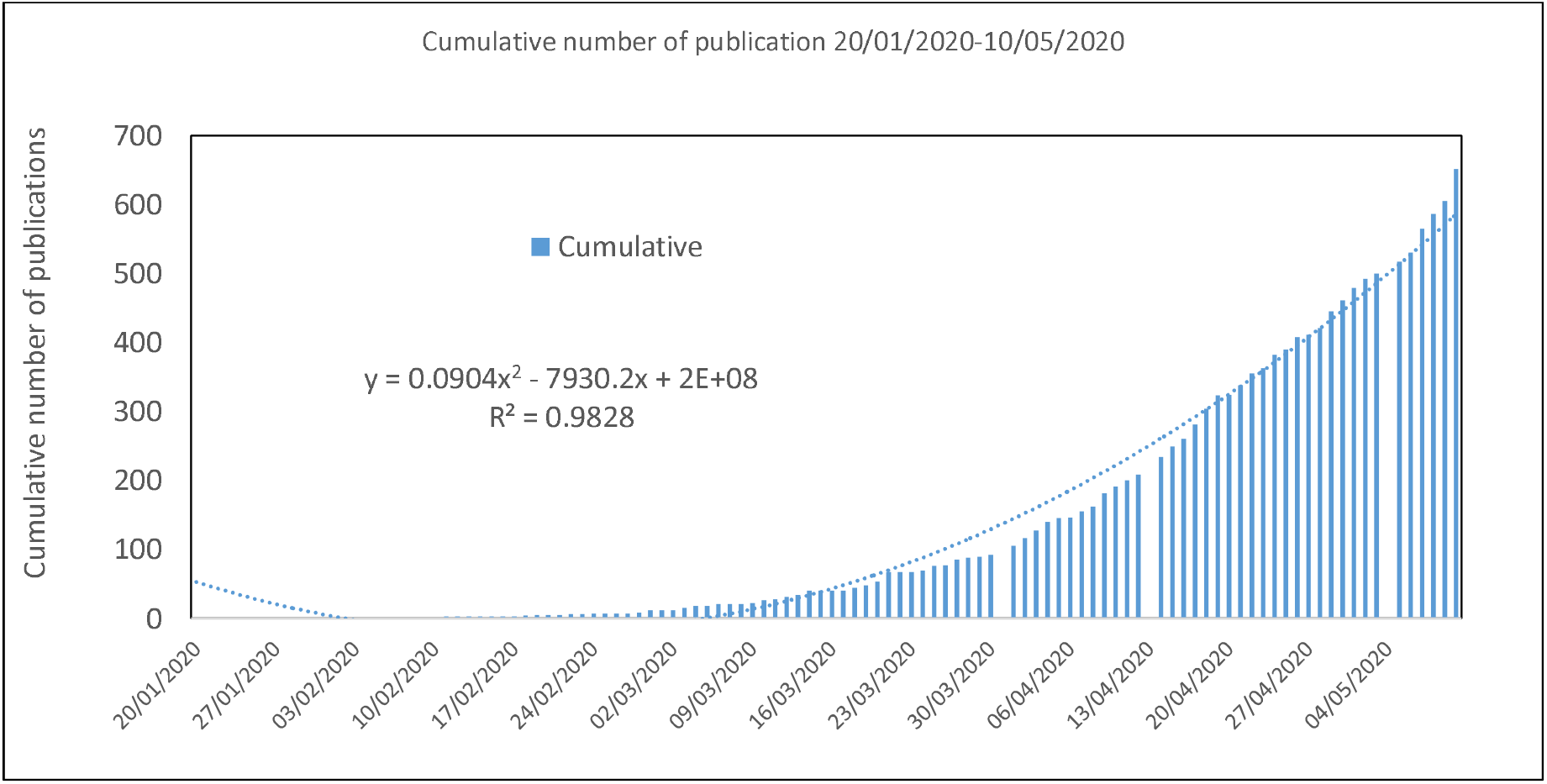
Cumulative number of publications as a function of days, PubMed All articles 20-02-2020 to 10-05-2020

**Figure-2b:**
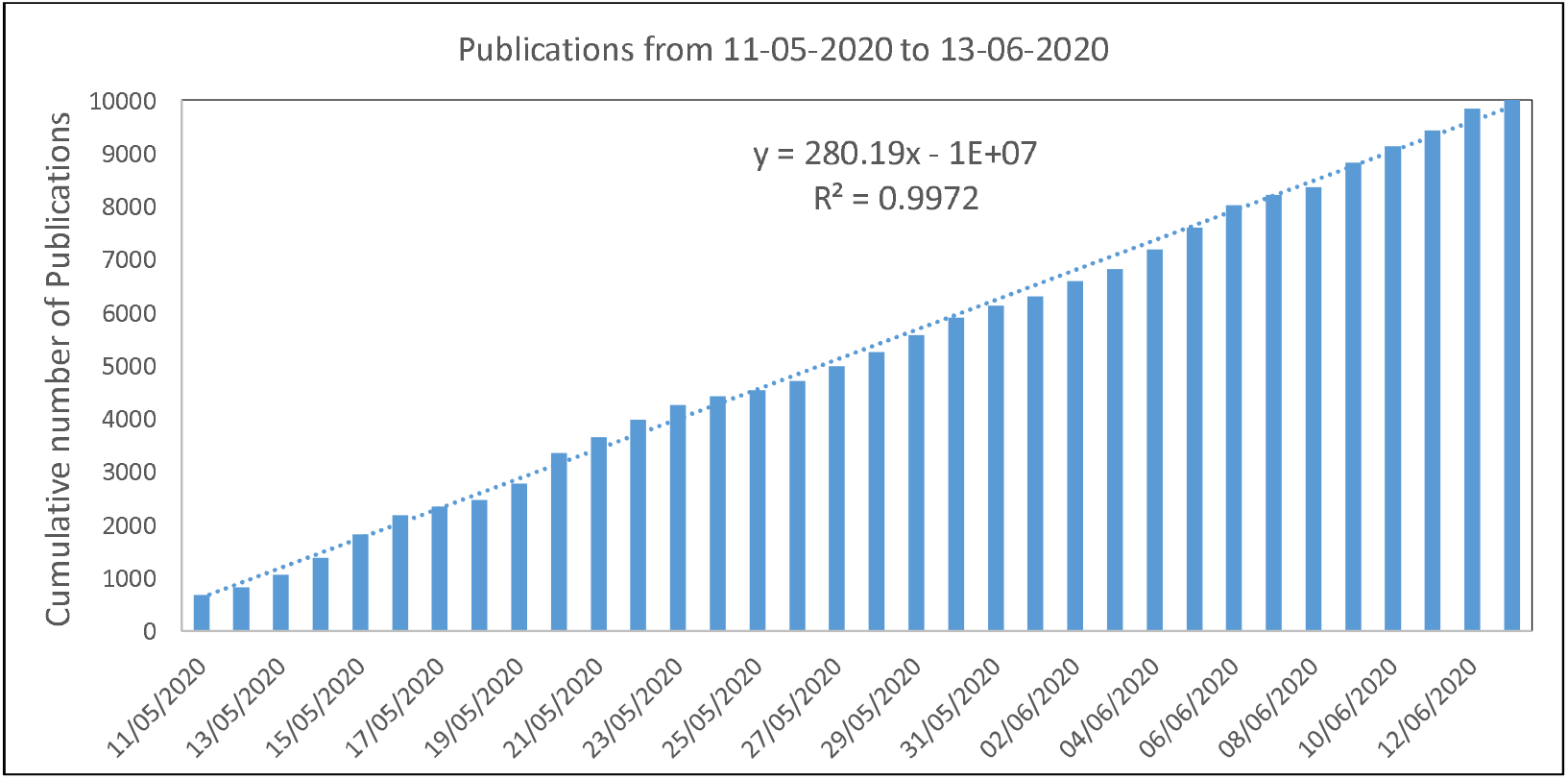
Cumulative number of publications as a function of days, Pubmed All articles 11-05-2020 to 13-06-2020

### PubMed: “all articles”: Number of publications vs. journals

The analysis was limited to 10000 articles published in 1868 different journals. Number of articles per journal was <1|15.2±10.3|191> (Median 2). While 750 journals published a single article, 695 journals published 2 to 5 articles (2336 articles in total) and the rest 423 journals published 6914 articles. The highest number of articles (191) was published by British Medical Journal (BMJ), followed by Journal of Medical Virology (138) and Dermatologic Therapy (113). Figures 3a and 3b show the number of publications as a function of the journal for range 1-9 and greater than or equal to 10 respectively. Appendix-1 shows the list of journals and their published articles.

**Figure 3a:**
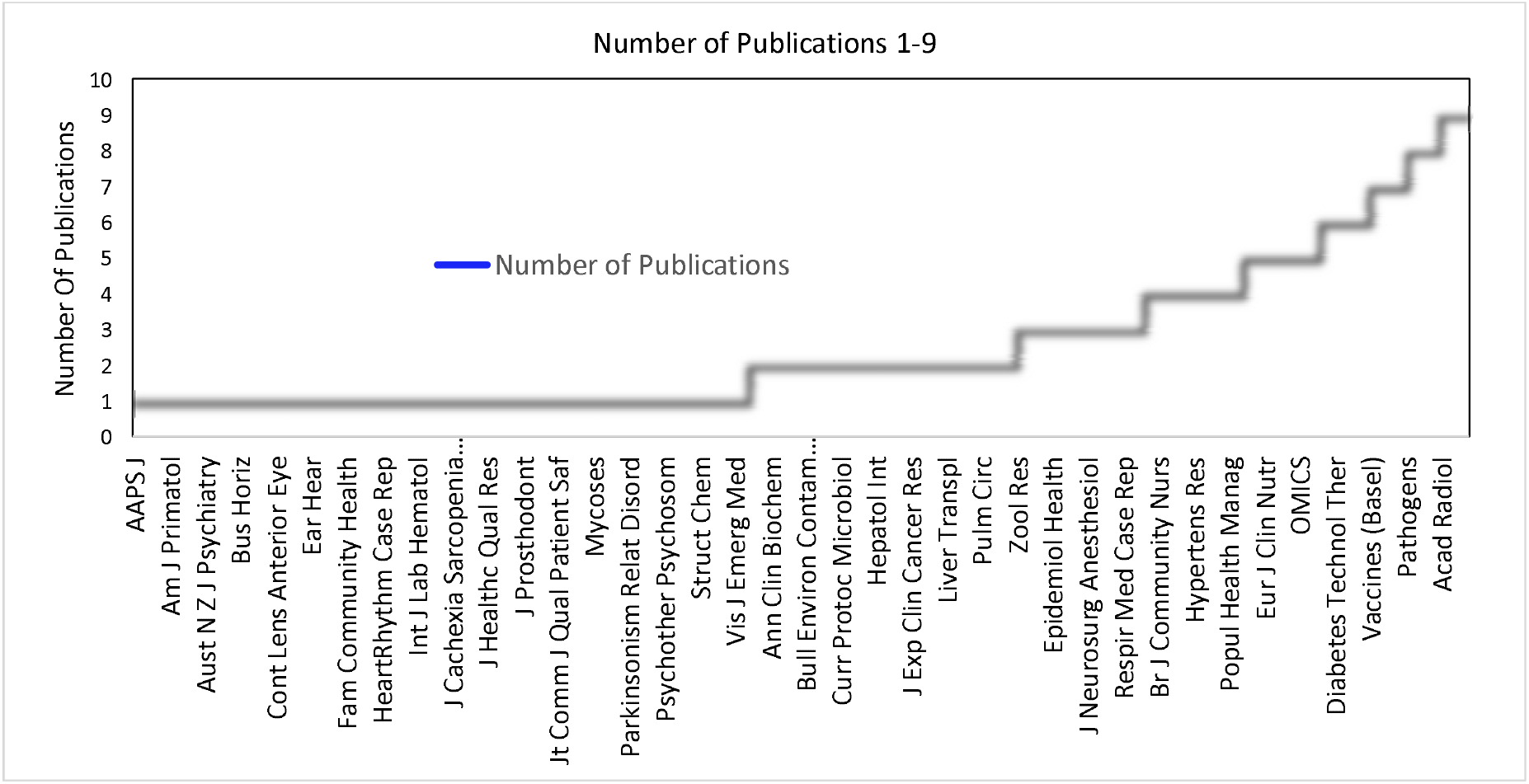
PubMed “all articles” Number of publications vs. Journal (For number of publications/journal ranging between 1 to 9) List of journals: supplementary data-1

**Figure 3b:**
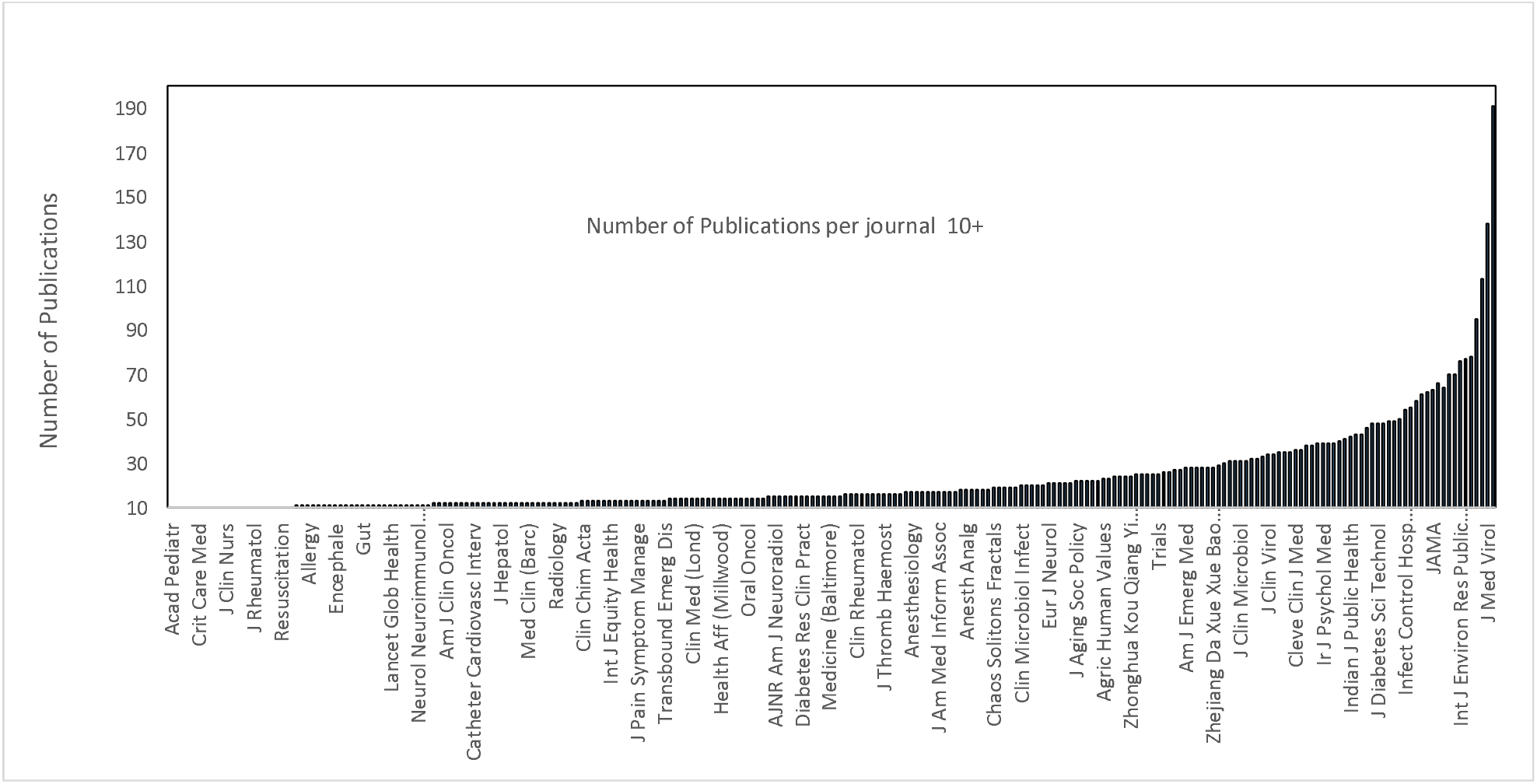
Pubmed “all articles” Number of publications vs. Journal (For number of publications/journal ≥10) List of journals: supplementary data-1

### PubMed: “all articles”: Number of publications vs. author

Figure-4 below presents the authors histogram. The average number of publications per author and the range was <1|1.2±1.4|71>. Median number of publications was 1. A total of 41083 authors were identified with 1 publication each, 3847 authors with 2 publications, 1583 with 3-10 publications, and 121 authors with 11-71 publications. A total of 8 authors have more than 50 publications. Top 20 authors representing 935 articles are of Chinese descent. Figure-4 presents the number of publications as a function of number of authors per publication. Authors’ names are provided in Appendix-2.

**Figure 4:**
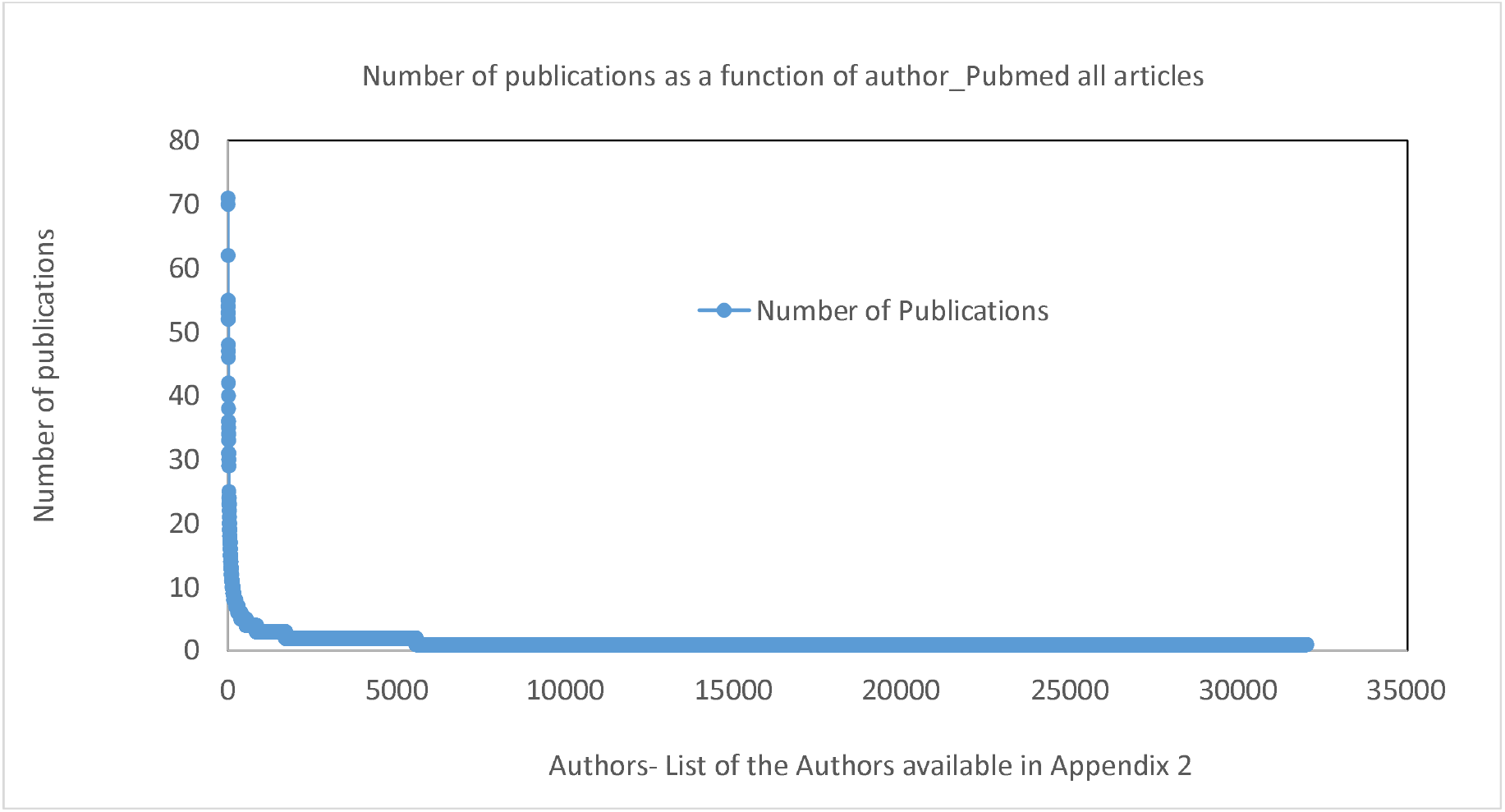
Authors vs. number of publications (PubMed “all articles”) - List of Authors available in supplementary data-3

### PubMed: Doubling time for the number of publications

The doubling time for the number of publications was calculated considering 3 publications on February 11, 2020 as the base. Between 11/02/2020-13/06/2020 (123 days) total of 12 doublings were observed (2^12^) and presented in figure-5. Mean doubling time was <2|10.3 ±4|15>.

**Figure-5:**
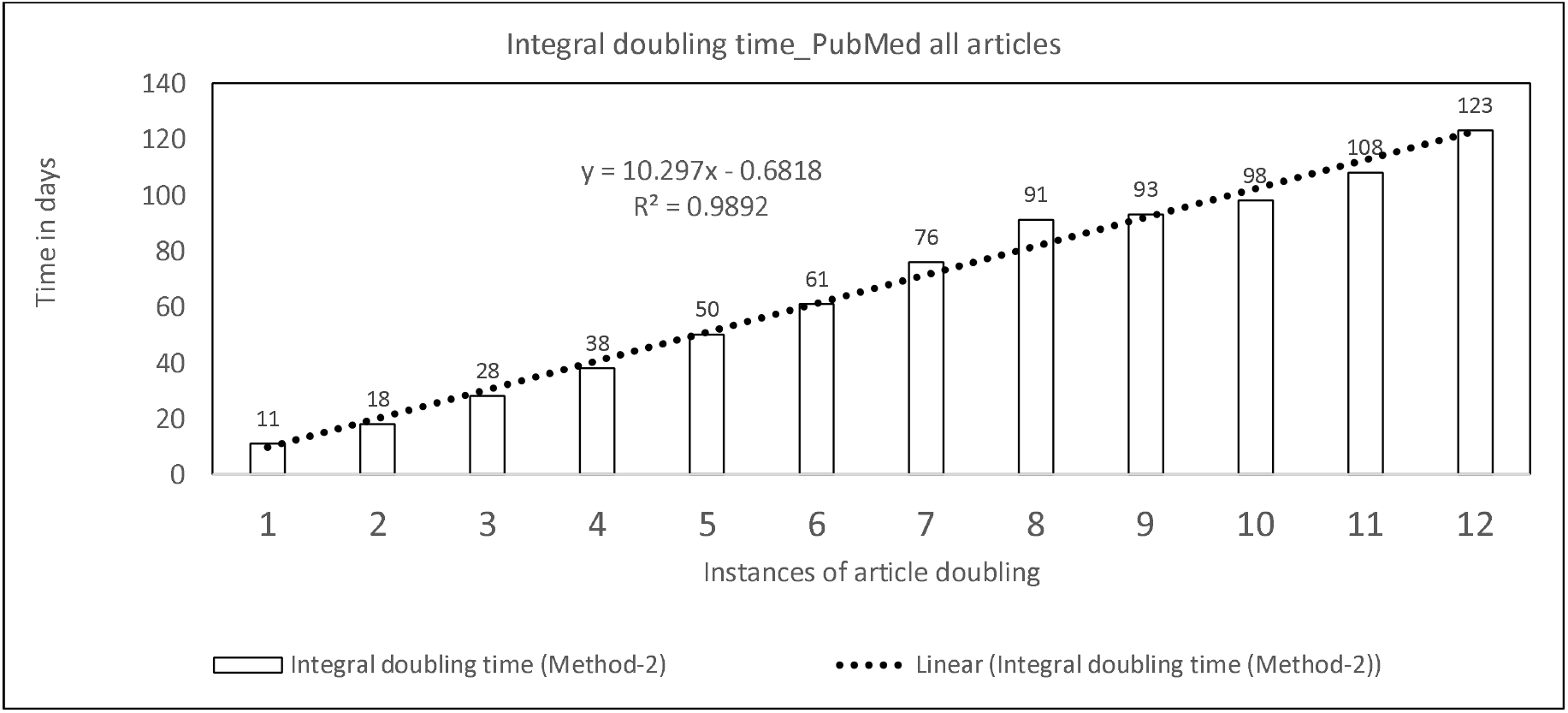
PubMed “all articles”: Article doubling as a function of days, total 12 doublings in the number of articles in 123 days

### 1b. PubMed: Review articles

Analysis of number of days taken for acceptance after submission (PubMed review article: Subgroup Analysis)

A total of 150 review articles that appeared in PubMed between 06/11/2019 to 28/04/2020 were analyzed for the number of days taken for acceptance (difference in days between acceptance and submission dates) (Fig-6). Three articles were accepted on the same date of submission, 18 in 1 day, 9 in 2 days, 5 in 3-5 days, 22 in 6-10 days, 41 in 11-20 days, 33 in 21-30 days. Only 19 articles were in review for more than 1 month. The average and median number of days in review were <1|17±17|77> days and 13.9 days respectively. If fitted with a straight line (y = −0.2144x + 9433, not shown) with the number of article review days reducing at a slope of −0.21 days (count)/day.

**Figure-6:**
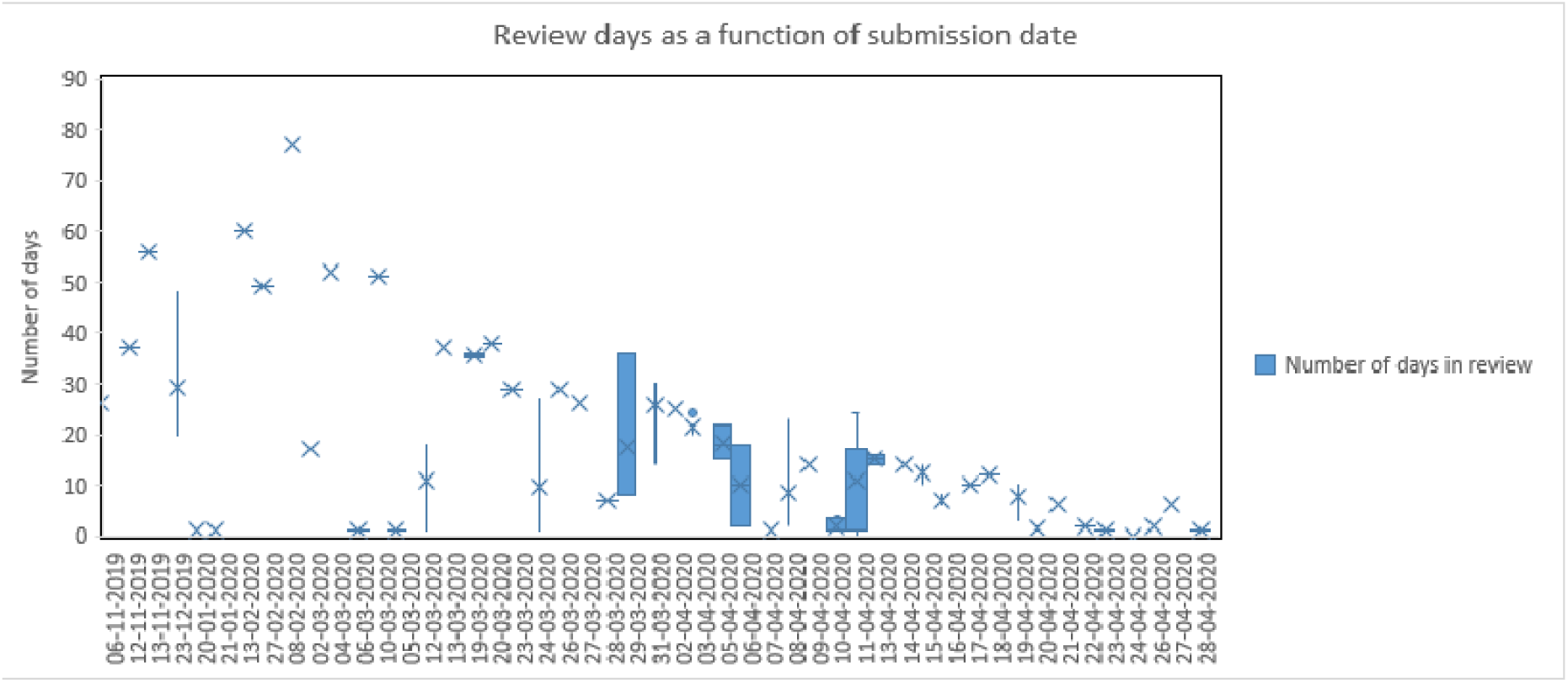
Analysis of days in review (acceptance-received date) days as a function of the submission date (PubMed review articles)

### 1C. Analysis of PubMed Clinical Trials

Between January-June 2020, PubMed showed a total of 17 randomized clinical trials (RCT) in (16 in English and 1 in Chinese language) published in 16 journals, with Lancet alone publishing 2 trials. However, after carefully reading each article, it was found that the publications from France and Italy were not RCTs and the total was reduced to 15: 13 articles from China and 2 from Brazil. A total of 300 independent authors were involved in these articles, the median number of authors per article is 13 and the range is from 6 to 65. Of these, only 6 articles have information on submission and accepted dates. The average and median review time was <0|10.7±15.3|41> days, 5.5 days respectively. Two review articles were from the same research group, Hainan General Hospital, Haikou, China which were accepted in 0 and 2 days respectively in a single journal (Complementary therapies in clinical practice). [9–10]

### 2. Elsevier

#### Publication Vs Days

Elsevier had published 5898 articles (excluding Erratum) for the observation period between 30/12/2019 to 25/05/2020. The average number of publications/day and the range were <1|77.6±125.3|767>, the median number of publications/day=27. Figure-7 shows the cumulative publication (in log scale) and the differential number of publications (in linear scale) as a function of days.

**Figure-7:**
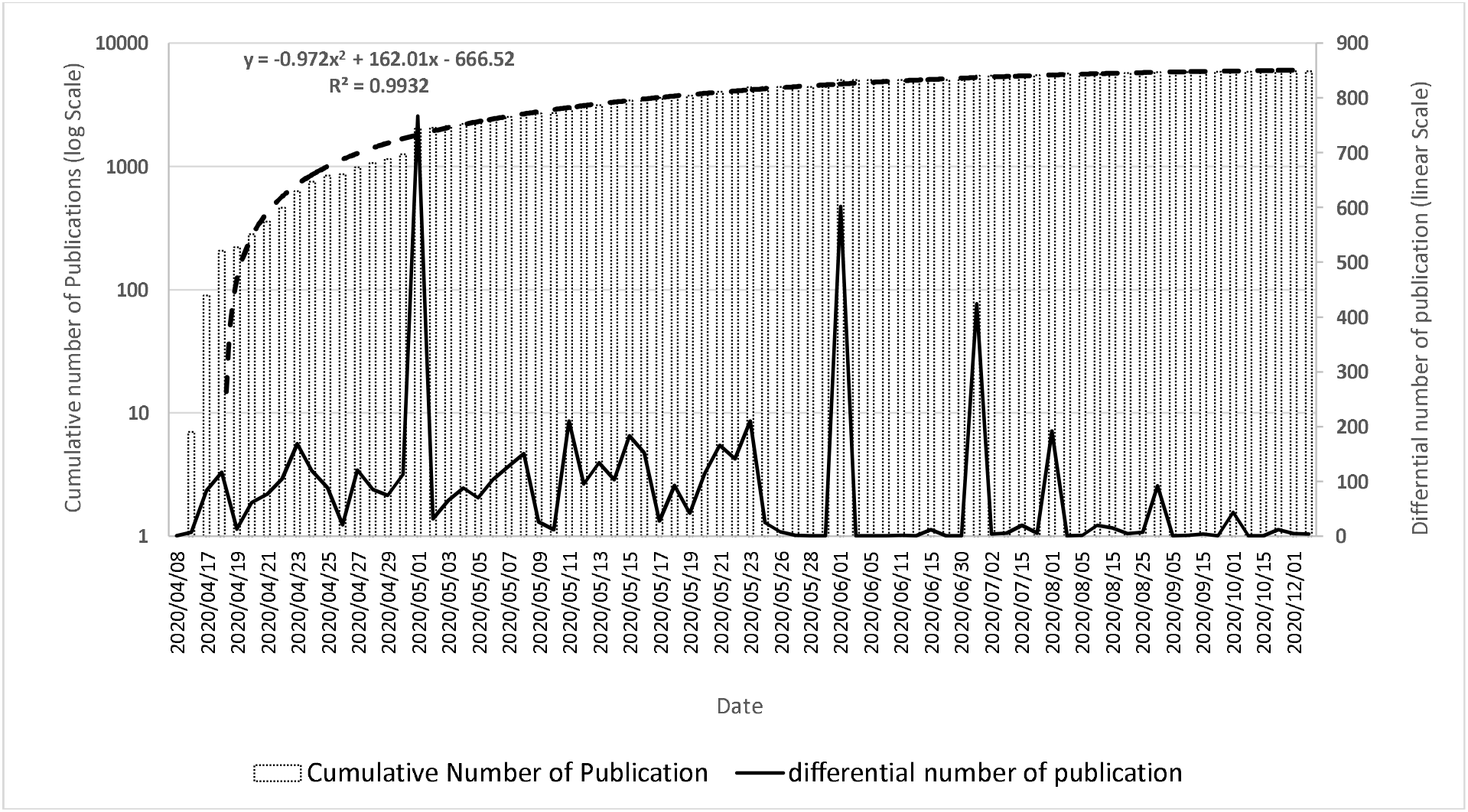
Elsevier: Cumulative and differential number of publications as a function of date

The cumulative number of publications as a function of date from Elsevier publishing house follows a parabolic trend with fitting equation as y = −0.972x^2^ + 162.01x - 666.52 and a fitting accuracy of 99.3%.

#### Doubling time

The cumulative number of publications in Elsevier encountered 13 doublings in number, with an average doubling time of 20.6 days. Figure 8 shows the cumulative number of articles as a function of doubling time, fits with exponential growth y = 0.7198e^0.6931x^.

**Figure-8:**
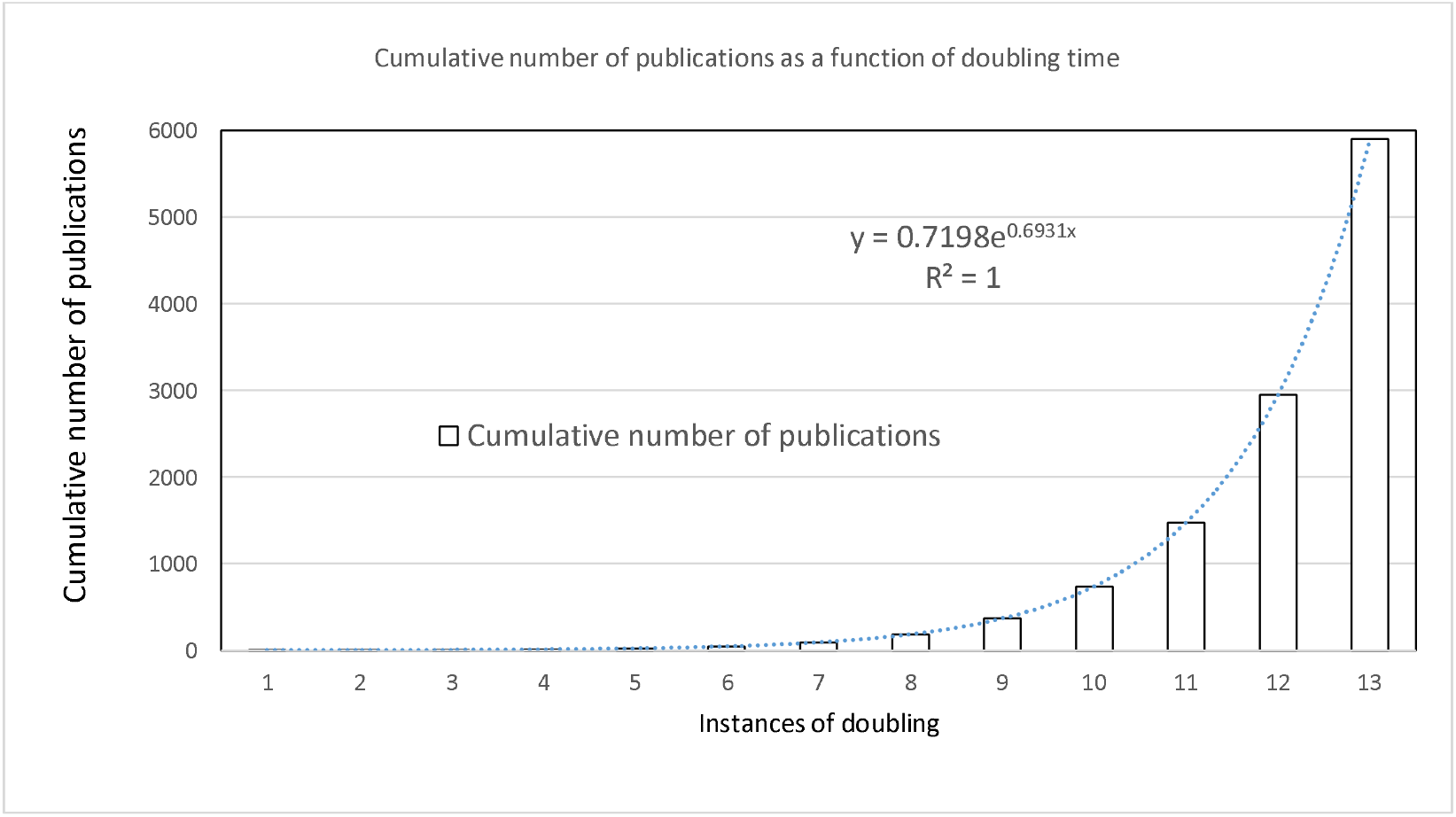
Cumulative number of publications as a function of doubling time (Elsevier)

#### Number of publications vs authors

Elsevier identifies a total of 27845 authors, with 22675 authors with lone articles, 3849 authors with 2 articles, 1297 authors with 3-10 articles and 23 authors with 10+ articles. The maximum number of articles for single authors was 34. Mean and range of the article is <1|12.3±0.9|34>, median =1 article. Figure 9 shows the number of publications as a function of the author (Elsevier).

**Figure 9:**
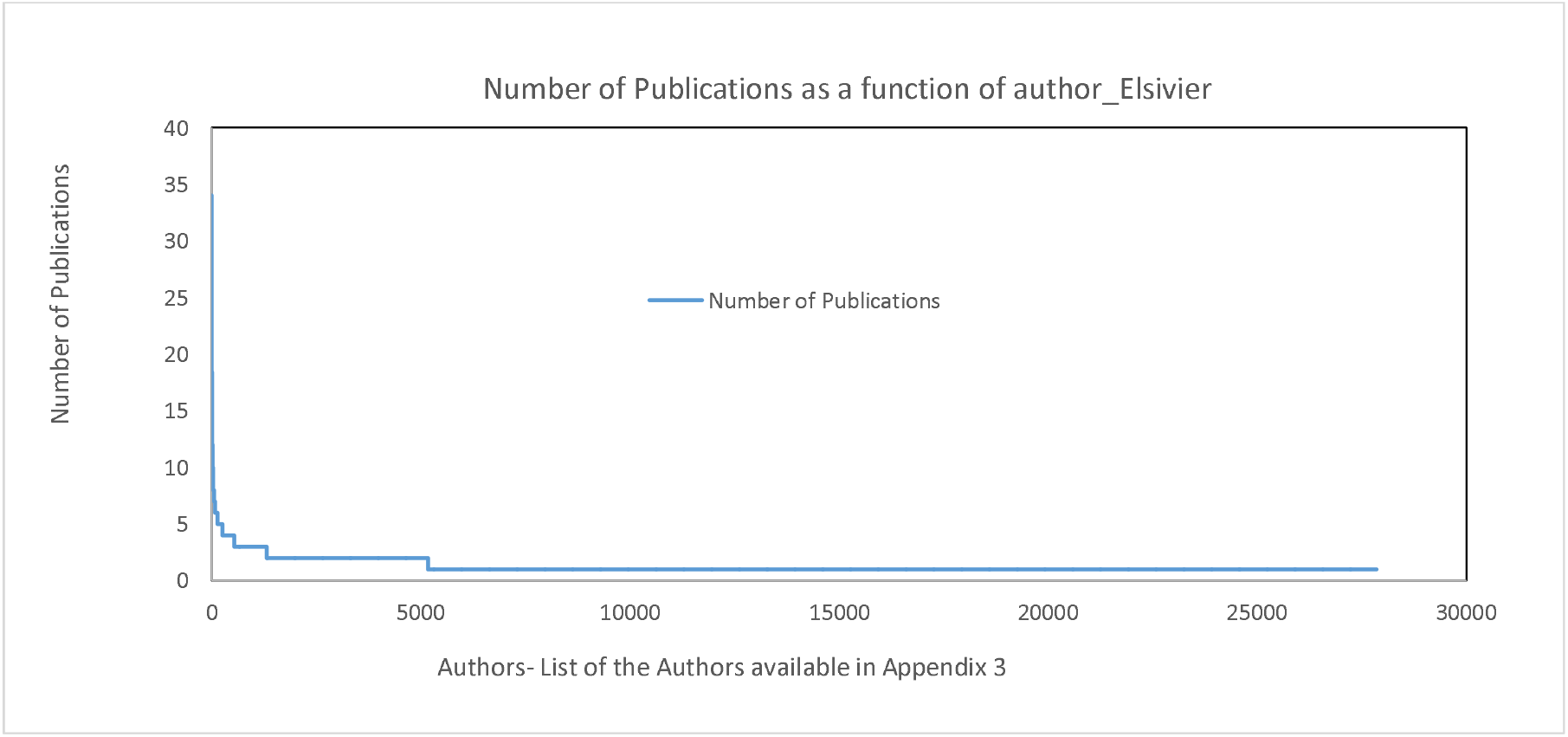
Number of publications as a function of author (Elsevier). Authors list available in supplementary data-3

### 3. Research Gate (RG)

Between 15/04/2020 and 30/04/2020 total 5395 documents related to COVID-19 was uploaded to RG. After carefully scrutinizing all entries, eliminating repetitions, erratum, presentations and comments, the number of useful documents reduced to 4180. A total of 1986 documents have full text.

#### Publication vs days

Figure-10 shows number of publications for each day during the last half of April 2020. Number of publication as a function of days shows a parabolic relationship. The average and median number publications were <40|258.4±130.8|441>/day and 273/day respectively.

**Figure-10:**
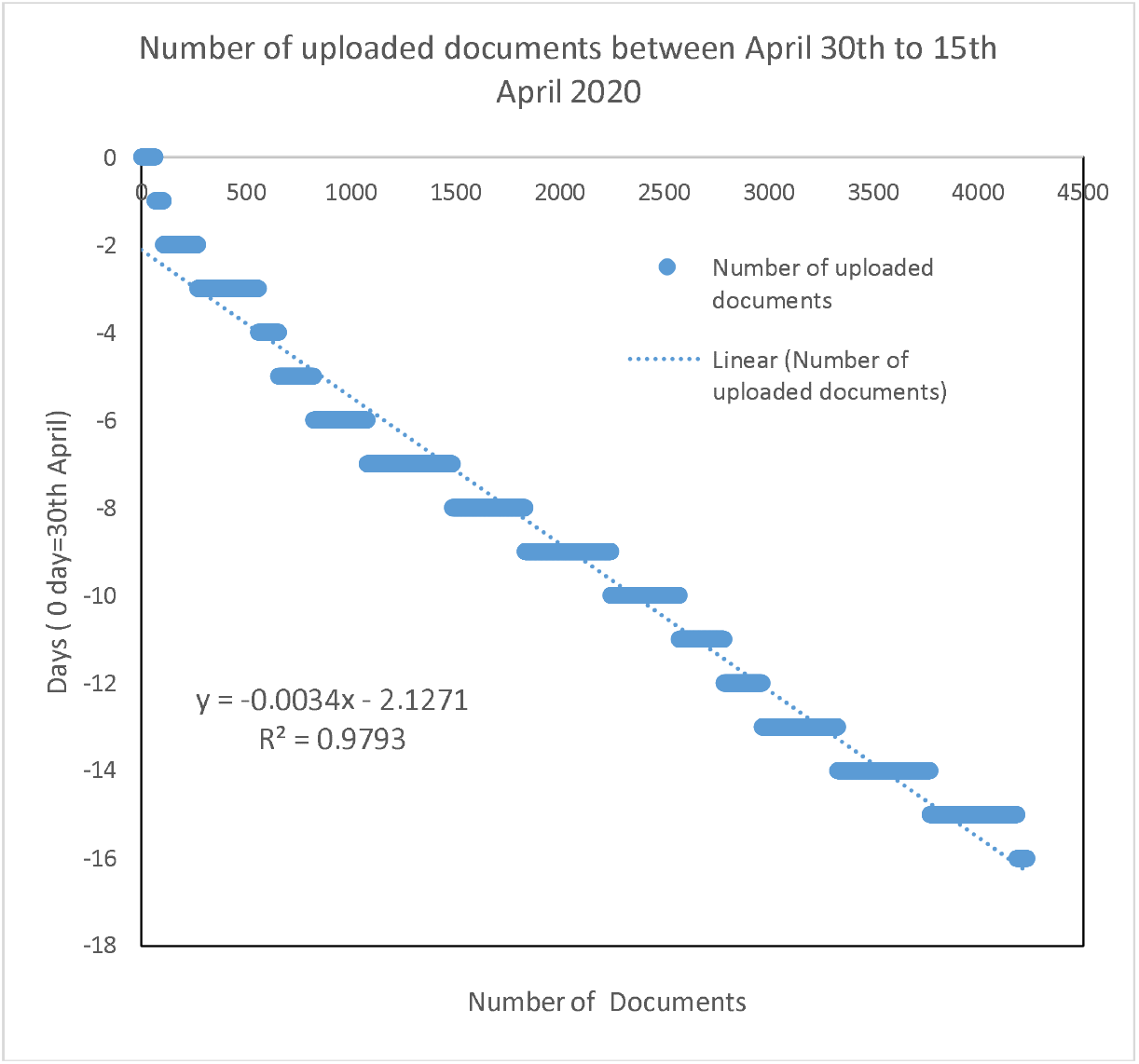
shows number in publication Vs Date in last two weeks of the April 2020 (Research Gate)

The documents related to basic science, diagnosis, drug and vaccine development, social and economic impact, public health, and treatment (with multiple option) were 635, 1622, 743, 927, 2179, and 1318 items respectively.

#### Doubling time

During the latter half of April, RG encountered 6 doublings in the number of uploaded documents. The doubling time for the number of publications is <1|2.3±2.0|6> days.

#### Number of publications vs. authors

Figure-11 shows the number of documents against the authors, with the full author list available in Appendix-4. A total of 3537 authors in RG have single article, whereas 257 people have 2, 18 people have 3, and another 18 people have 4 to 9 documents.

**Fig 11:**
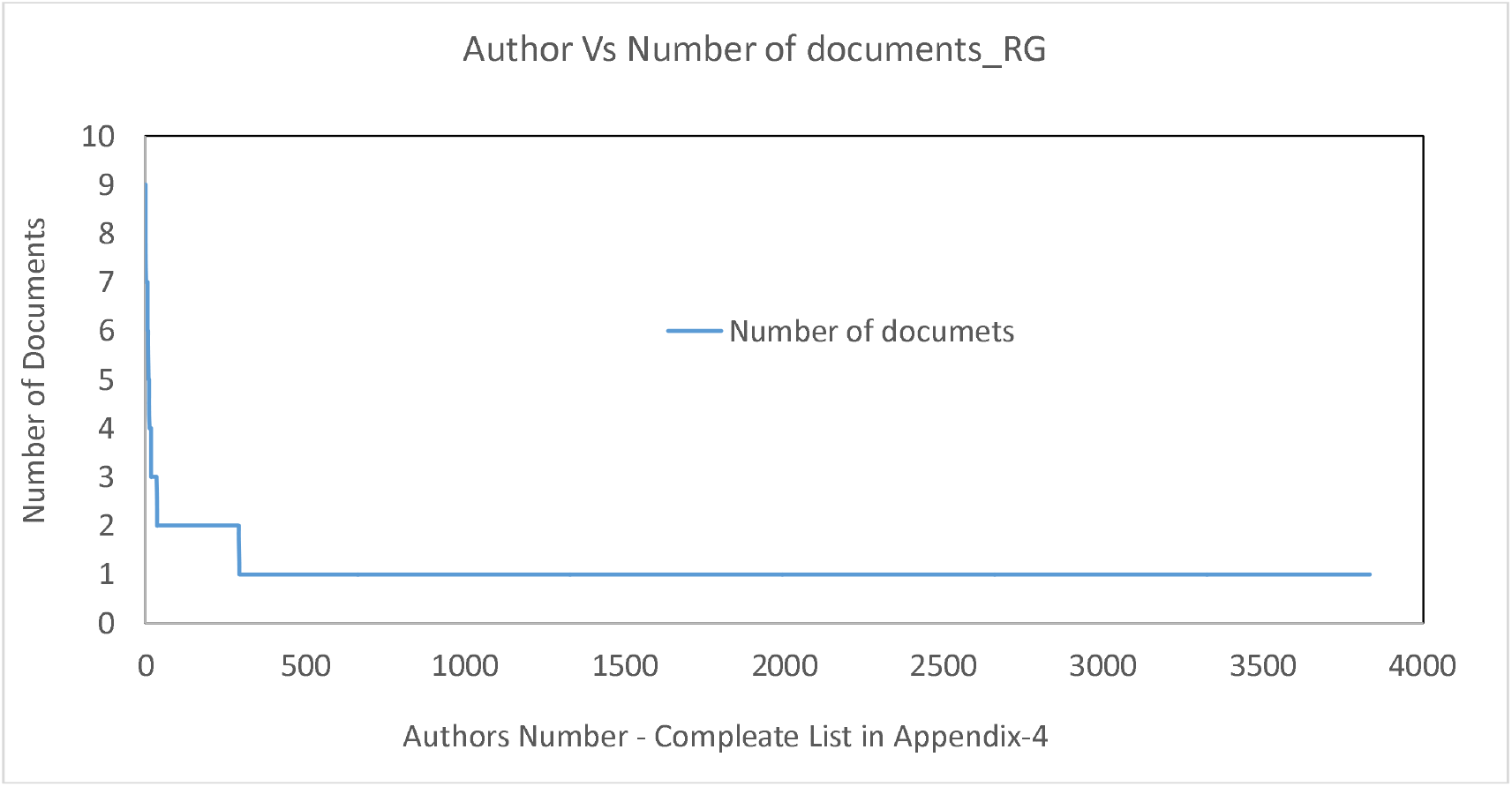
Author vs number of documents uploaded in Research Gate (complete author list available in Supplementary data-4)

## Discussion

Coronavirus disease 2019 (COVID-19) has stimulated the global scientific community to take unprecedented interest on a single subject aimed at learning more about the disease, sharing knowledge immediately, and undertaking concerted, evidence-based efforts to manage the situation. Calls for ensuring that all research findings relevant to COVID-19 be made available openly and promptly had begun to be issued as early as in January. In March, UNESCO mobilized countries to promote open science and data sharing to manage this crisis. Simultaneously, a global research roadmap for COVID-19 was issued by the World Health Organization (WHO) which pointed to the existing knowledge gaps. It has also drawn up timelines for the implementation of specific research actions.

A large number of journals have made the Covid-19 literature free to access. With the rapid increase in the number of publications, some of the journals have already crossed their December 2021 issue, and many journals have filled until the December 2021 issue (as on 12-06-2020).

In our study, we found the growth of articles as a function of time in different databases to be exponential, parabolic with low “a” value and linear with a very high gradient, pointing to a sudden, steep growth in number of articles. The doubling times in different databases were also varying fast and reducing continuously.

The situation calls for addressing some pertinent questions: (1) How is it possible to bring out such a large volume of literature in such a short time? (2) What is the reliability of the data collection, analysis, and literature review? (3) When articles, including randomized trials, are being accepted in a very short time, some showing same day acceptance, what is the quality and reliability of the review? (4) How will it be possible to read and comprehend this large volume of literature to extract useful information? (5) Whether this plethora of literature is actually translating or will it ever translate into any medical, social, or economical benefit? (6) How much of the information is real progress and how much is mere repetition?

Probably the answer for most of the above questions shall be in negative. It is interesting to note that 8 authors have more than 50 publications each in the analyzed time. The growth pattern of the number of literature is also alarming: doubling every 10.3 days in PubMed with a daily increase of 282.5±110.3 articles. Another matter of concern is the large presence of pseudoscientific facts, that are potentially harmful, in these publications. [12] This tendency of somehow to get an article published, is seen across the board, from established researchers of most reputed medical schools to undergraduate scholars from little known universities. It is therefore not surprising that reputed journals had to do one of the biggest ever retraction of scientific papers in the modern history, [13] marking a grim testimony to the chinks in the peer review process and robustness of audits. Even more concerning is that the authors of these articles are well established researchers. [14–15]

These retracted papers are most often fabricated and serve as examples of how the Covid-19 has overwhelmed the peer review process with a huge rush for publications. [16] Making use of the prevailing situation, some new predator journals have also popped up, usually paid journals, to present these pseudoscientific facts. [17]

Nevertheless, retraction of a paper does not necessarily cease completely its use or citation. Several examples can be found in which retracted papers are cited several months after retraction. [18]

Our analysis also shows that authors from Peoples Republic of China (PRC) are the leading contributors in the Covid-19 literature. First 20 leading authors in PubMed contributing an average of approximately 50 articles each, are of Chinese origin. A total of 87% of the controlled trials in PubMed has originated from PRC; 90% investigators in controlled trials are of Chinese descent.

The other alarming tendency among the journals and reviewers are very short review time, which is decreasing at a rate of −0.21 days with every passing day. Articles are often accepted in the same day or within a week. In PubMed RCT section we found two randomized controlled trials which were accepted in 1 day.

Another interesting feature in our analysis is the tendency of attaching the articles with “Covid-19”. [19] This is perhaps being done to increase the chance of publication, but is potentially a concerning tendency.

We have presented the data from a closed data sharing platform PubMed operated by National Institute of Health with no role of end users, data from one publishing house (Elsevier) and an open data sharing platform Research Gate operated by the end users. This helps in truly reflecting the overall scenario of all kinds of closed and open scientific data sharing platforms on Covid-19 data dynamics.

Has the interest in publishing more on Covid-19 has taken undue precedence over other subjects? The answer to this pertinent question can be judged from the fact that the number of publications on Covid-19 stands at 15354 that far exceeds the number of publications on cancer (in its all derivatives) which is a paltry 4718 (as on 14/07/2020).

## Conclusion

Our analysis of the dynamics of publications as a function of days, journals, authors in three different types of databases has undoubtedly proved the skewed bias toward excessive, unwarranted publishing on Covid-19 pandemic. Though the disease has been declared as a global pandemic, it does not warrant unwanted literature piling up continuously. Research groups, publishing houses, journals, reviewers and all associates need to realize that the race for publications related to Covid-19 needs to be pragmatic, not a blind one. Publishing articles based on scientifically unimportant, pseudoscientific, fabricated, unreliable and harmful facts just to increase the number of publications and citations is actually harmful to society and disservice to the scientific community.

